# Songbird mesostriatal dopamine pathways are spatially segregated before the onset of vocal learning

**DOI:** 10.1101/2023.05.18.541314

**Authors:** Malavika Ramarao, Andrea Roeser, Caleb Jones, Jesse H. Goldberg

**Affiliations:** Department of Neurobiology and Behavior, Cornell University, Ithaca, NY 14853, USA

## Abstract

Diverse dopamine (DA) pathways send distinct reinforcement signals to different striatal regions. In adult songbirds, a DA pathway from the ventral tegmental area (VTA) to Area X, the striatal nucleus of the song system, carries singing-related performance error signals important for learning. Meanwhile, a parallel DA pathway to a medial striatal area (MST) arises from a distinct group of neighboring DA neurons that lack connectivity to song circuits and do not encode song error. To test if the structural and functional segregation of these two pathways depends on singing experience, we carried out anatomical studies early in development before the onset of song learning. We find that distinct VTA neurons project to either Area X or MST in juvenile birds before the onset of substantial vocal practice. Quantitative comparisons of early juveniles (30-35 dph), late juveniles (60-65 dph), and adult (>90 dph) brains revealed an outsized expansion in the number of Area X-projecting and MST-projecting VTA neurons over development. These results show that a mesostriatal DA system dedicated to social communication can exist and be spatially segregated before the onset of vocal practice and associated sensorimotor experience.

## Introduction

Midbrain dopamine (DA) neurons project to diverse striatal areas and contribute to reinforcement learning by signaling reward prediction error, the difference between actual and predicted reward [1]. In classic models, this DA reinforcement signal is posited to be globally broadcast throughout the brain so that an animal can learn arbitrary stimulus-response associations[2-4]. Closer examination of the pathway-specific DA signaling has shown that DA signals can vary across distinct striatal regions [5-8]. For example, medial, lateral, dorsal and ventral zones of the mesostriatal projection can exhibit distinct responses to movements and errors in reward prediction[9]. The DA projections to the posterior tail of the striatum and specific accumbens regions may even exhibit non-canonical responses to aversive stimuli [10, 11]. These studies raise the possibility that mesostriatal DA systems do not always broadcast a global scalar reinforcement signal but rather may provide a vector of evaluative feedback that targets specific parts of the animal’s motor system to produce optimal policies in complex environments [12].

The evidence that distinct anatomical channels of the mesostriatal system exist and may exhibit distinct aspects of evaluative feedback arises primarily from studies in adult animals. This raises a critical question: do distinct mesostriatal DA systems achieve their segregation by synaptic pruning early in life, possibly influenced by experience? If this is the case, then juvenile animals may exhibit less anatomical segregation than adults. Alternatively, segregation might be specified genetically and be evident in juveniles before the onset of learning.

The songbird provides an ideal model system to examine mesostriatal pathway development. First, songbirds have a discrete neural circuit, the song system, that is dedicated to song learning [13]. Importantly, the song system contains a specific DA projection from VTA to Area X, the singing-specialized striatum [14, 15]. Past studies in adults showed that neurons in VTA that project to Area X (VTA_X_ neurons) exhibit performance evaluation signals important for learning [16-20]. In addition, VTA_X_ neurons are distinct from VTA neurons that project to a more medial striatal region (MST) unassociated with vocal circuits [14, 15]. We have been perplexed by two aspects of the VTA-Area X and VTA-MST pathways in songbirds. First, even though VTA_X_ neurons and MST projecting VTA neurons (VTA_MST_ neurons) arborize in distinct striatal regions, they can be spatially intermingled, with cell bodies adjacent to one another [15]. Second, VTA_X_ neurons carry singing related auditory error signals, but neighboring ones that project to MST do not. In electrophysiological recordings, antidromically identified VTA_X_ neurons that exhibited error signals during singing were often recorded in the same electrode penetration, and at the same site, as VTA neurons that did not project to Area X and did not encode performance error [17, 18]. How do the neurons in VTA that encode auditory error ‘know’ to send their axons to Area X and not to MST? One possibility is that a single, global mesostriatal DA system exists early in development, but that the act of singing contributes to an experience-dependent process that causes separate pathways to form. Singing-related auditory signaling found in some VTA neurons promotes synapse formation with singing-related neurons in Area X, analogous to how early visual experience enables normal visual system development and, for example, ocular dominance columns [21]. Alternatively, the VTA_X_ projection could be genetically specified, for example, by cell-class specific markers that control axon guidance [22].

Zebra finches learn to sing through a gradual process of trial and error. In early development, until about 35 days post hatch (dph), young birds hear a tutor song and form an auditory memory, or an internal model, to be imitated [23]. Between 35-45 dph, juvenile male zebra finches begin to sing subsong, highly variable vocalizations akin to human vocal babbling [24]. Over the next 50 days, the song gradually becomes more stereotyped and similar to the tutor song, finally crystalizing by 90-100 dph [25, 26]. DA-based reinforcement learning is known to involve plasticity in mammalian cortico-striatal synapses [27], and DA-modulated plasticity of cortical inputs to Area X, measured in brain slice recordings, is observable at intermediate stages of vocal development (47-52 dph)[28, 29], suggesting early innervation of Area X by DA neurons. Importantly, topographic connectivity of cortico-basal ganglia pathways during song system development is affected by manipulations of auditory experience early in life, suggesting that experience can play a role in refining vocal circuitry [30, 31].

This vocal learning process provides a clear timeline for testing the development of the mesostriatal DA systems [32]. By injecting distinctly colored retrograde neuronal tracers into Area X and MST in juvenile birds and examining tissue days later, we tested if VTA cell bodies projected to Area X, MST, or both regions, in the brains of birds that prior to subsong singing (early juveniles, 30-35 dph) and birds that were in the process of learning their distinct song motif (late juveniles, 60-65 dph). By repeating these experiments in adult birds, we tested for the replicability of past studies and compared cell counts in juveniles and adults. If the dopaminergic projections from VTA to MST differentiate based on singing experience, as has been shown in forebrain nuclei of the song system[30], then prior to song learning, single neurons in VTA may project to both MST and Area X. On the other hand, if DA projections from VTA to MST are pre-determined, for example by genetic influences, then VTA neurons will already be segregated by projection area in juveniles. We discovered that the two pathways were completely segregated early in development, supporting the second outcome where a song system-associated DA population exists even before the onset of vocal practice.

## Materials and Methods

### Animals

All animal procedures were in accordance with NIH guidelines, the Institutional Animal Care and Use Committee of Cornell University (ID# 2018-0026), and the New York State Department of Health. We used data from 26 healthy male zebra finches (Taeniopygia guttata), age range 30–700 days post-hatch (dph). Birds were obtained from our breeding colonies in Ithaca, New York. Prior to surgery, birds at least 50 dph, were in constant social contact with other males in large colony cages and had visual and auditory contact with females. Birds below 50 dph were in constant social contact with both male and female zebra finches in a large breeding cage. Loosely attached leg bands were used as identifiers for each bird after at least 50 dph. Post-surgery, birds were in individual cages, but had visual and auditory contact with both males and females in separate cages. Perfusion was performed 5 days after surgery. Pre- and post-surgical birds were located in rooms with controlled temperature and humidity conditions on a 12/12 hour day/night schedule with constant access to food and water (seed mix and water refreshed daily by animal technicians employed by Cornell University). Environmental enrichment consisted of mirrors and perches of diverse sizes, and cuttlebones in cages. Weekly enrichment consisted of vegetables, egg, and water baths.

For surgeries, animals were anesthetized with 1-2% isoflurane, and anesthesia was assessed every 5 minutes and isoflurane levels were adjusted (between 1-3%) accordingly. The scalp is cleaned with Betadine (10% in distilled water). After craniotomies were made in the correct locations, 30 nL of fluorescently labeled cholera toxin subunit B (CTB Alexa 555, Molecular Probes) was injected bilaterally into Area X and 30 nL CTB Alexa 647 (Molecular Probes) was injected into MST bilaterally at the coordinates shown in Table 1. 10 minutes prior to the end of surgery, buprenorphine and enrosite were injected, and a mixture of 5% lidocaine cream and bacitracin antibiotic ointment was applied to the scalp. Following surgery, animals were placed in a small holding cage, recovered quickly, and were typically able to eat and drink within 15 minutes. After recovery, the animals were assessed approximately every hour for 4-6 hours and then assessed at least 3 times a day for 2 days after the surgery. If post-op recovery did not appear normal (ex. Bird does not recover from anesthesia, regain balance), then the animal was immediately euthanized.

**Table 1:**
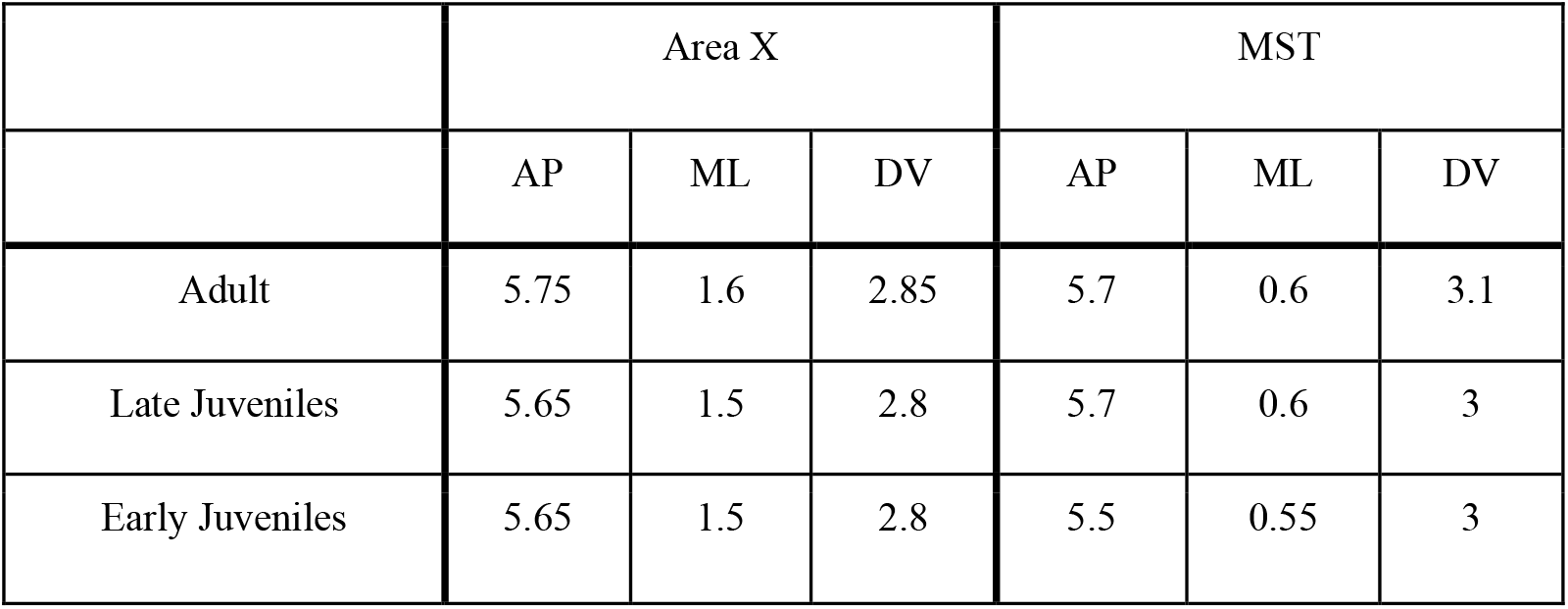
Stereotaxic injection coordinates for all surgeries performed on the zebra finches. All injections were carried out at a 20° head angle, coordinates zeroed at the lambda bifurcation, and performed bilaterally.

|  | Area X |  |  | MST |  |  |
| --- | --- | --- | --- | --- | --- | --- |
|  | AP | ML | DV | AP | ML | DV |
| Adult | 5.75 | 1.6 | 2.85 | 5.7 | 0.6 | 3.1 |
| Late Juveniles | 5.65 | 1.5 | 2.8 | 5.7 | 0.6 | 3 |
| Early Juveniles | 5.65 | 1.5 | 2.8 | 5.5 | 0.55 | 3 |

Surgeries were performed on three age groups, adult birds (>90 dph, n=12 hemispheres, n=6 birds), late juveniles (60-65 dph, n=7 hemispheres, n=5 birds), and early juveniles (30-35 dph, n=7 hemispheres, n=4 birds).

### Perfusion and Histological Processing

Injected birds were perfused 5 days post-injection with 4% paraformaldehyde diluted in phosphate-buffered saline (PBS) and stored at 4°C for 24-48 hours. Brains were then sliced using a Leica VT1000 S vibratome into 100 *μ*m sagittal sections in PBS, mounted under Polyvinyl alcohol mounting medium with DABCO (Sigma Aldrich), and stored at 4°C.

### Image Processing and Cell Counting

Slices were imaged using a Leica DFC345 FX fluorescent microscope, and pictures were taken of the injection sites, HVC, and all slices containing VTA. HVC was imaged to confirm Area X injection accuracy, since HVC projects to Area X in both juvenile and adult zebra finches [31, 33]. The number of cells in VTA labeled by the different fluorophores were manually counted and their precise locations were documented (Supplementary Table 1). Cell counting and co-labeling determination was performed manually using the images generated on the fluorescent microscope and ImageJ software (Fig 1C-H, Supplementary Fig 1).

**Figure 1:**
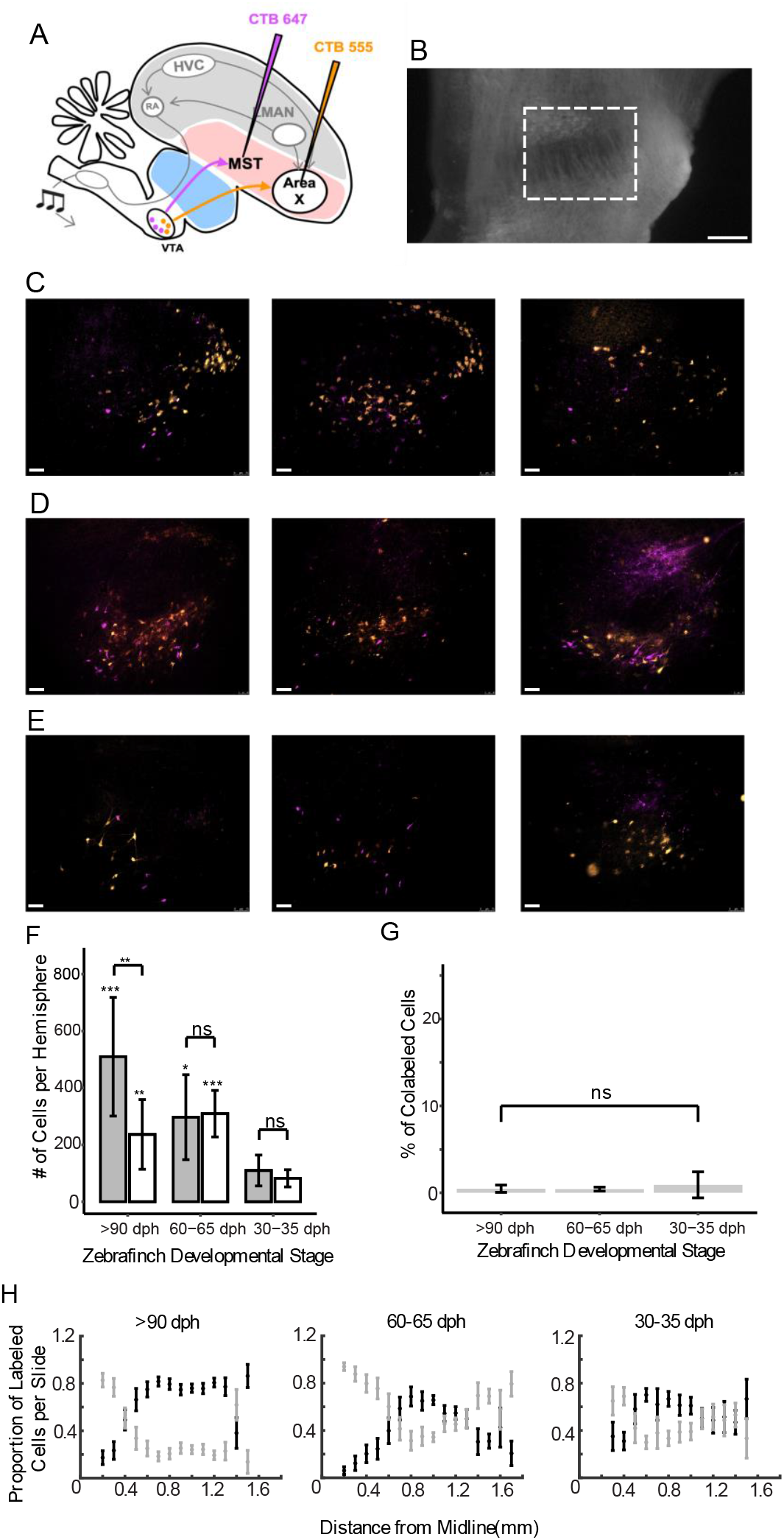
VTA Projects to Area X and MST. (A) Schematic of the experimental strategy used to label VTA projections to MST and Area X; (B) Darkfield image with expanded view of VTA, denoted by dashed white rectangle, scale bar 250 μm; (C) MST (magenta) and Area X (orange) projecting neurons in VTA in adults; (D) late juveniles 60-65 dph; (E) early juveniles 30-35 dph. Scale bar in C-E, 75 um. Dorsal is up; anterior is to the right. (F) Average cell counts per hemisphere of populations projecting to Area X (dark gray) and MST (white) from VTA in all three injection sites with an n of 12, 7, and 7 hemispheres respectively. Error bars reflect standard deviation of the mean. Asterisks above late juveniles and adult populations show results of two sample t-tests when comparing to appropriate early juvenile populations. Asterisks above bars show results of two sample t-tests between neuron populations within the same age group. (G) Percent of co-labeled cells in VTA that project to both MST and Area X in all three injection sites with an n of 12, 7, and 7 hemispheres respectively. Error bars reflect standard deviation of the mean. Significance score shows the results of two sample t-tests between all categories. Two sample t-tests were performed between all combinations of the three categories. (H) Proportion of Area X (black) and MST (gray) projecting cells over total labeled cells in the brain slice, progressing from medial to lateral. Error bars reflect standard error of the mean.

## Results

### VTA projections to MST or Area X are segregated before the onset of vocal practice

In a previous study using dual-color retrograde tracing experiments with injections into Area X and MST, the VTA_MST_ projection was found to be distinct from the VTA_X_ neuron population in adults, with a colabeling of < 5% [15]. To test if this segregation of DA pathways exists before the onset of vocal learning and examine changes over development, we repeated these experiments in both early juvenile animals (<35 dph, n=7 hemispheres), intermediate juveniles (60-65 dph, n=7 hemispheres), and adults (>90 dph, n=12 hemispheres) (Methods) (Fig 1, Supplemental Fig S1). We found that the percent of colabeled VTA_X_ and VTA_MST_ neurons was similarly negligible in adults and in both early and intermediate juvenile groups (11/1346 cells colabeled in 30-35 dph, n=7 hemispheres; 19/4247 cells colabeled in 60-65 dph, n=7 hemispheres; 37/8955 cells colabeled in >90 dph, n=12 hemispheres; p > 0.2 for comparisons between all three populations, 3 unpaired t-tests), with a colabeling of < 1% (Fig 1G). Thus the VTA_MST_ and VTA_X_ pathways are segregated even before the onset of substantial vocal practice.

While the percent colabeling between adults and juveniles was not significantly different, the topographical separation between the two populations widened as juveniles transitioned to adulthood. In early juveniles, the VTA_MST_ and VTA_X_ projections co-mingled in the anterior-dorsal region of VTA (Fig 1H). But in adults and late juveniles, the VTA_MST_ neurons, though still continuous and co-mingled with VTAx neurons, appeared to cluster in two distinct territories: one lateral to the VTA_X_ neurons in dorsal VTA and another medial to the VTA_X_ neurons in the ventral-posterior region of VTA (Fig 1H). These results are compatible with the idea that the VTA_X_ neuronal population expands during development and migrates into the region, displacing MST projectors medially and laterally.

### Outsized expansion of the VTAx population over development

To test the possibility of VTA_X_ population expansion during vocal learning, we examined how the number of VTA_X_ and VTA_MST_ neurons changed over development by comparing cell counts and locations in the juvenile and adult datasets. In early juveniles, the number of VTA_X_ and VTA_MST_ neurons was not significantly different (avg. 110 VTA_X_ neurons/hemisphere vs avg. 82 VTA_MST_ neurons/hemisphere; p value of 0.271; n=7 hemispheres; Fig 1F). In late juveniles, the number of VTA_X_ and VTA_MST_ neurons was not significantly different (avg. 297 VTA_X_ neurons/hemisphere vs avg. 309 VTA_MST_ neurons/hemisphere; p value of 0.851; n=7 hemispheres; Fig 1F). However, in adults the number of VTA_X_ neurons dramatically expanded relative to the number of VTA_MST_ neurons (avg. 510 VTA_X_ neurons /hemisphere vs avg. 237 VTA_MST_ neurons/hemisphere; p<0.05; n=12 hemispheres; Fig 1F), and the number of VTA_MST_ neurons in adults does not significantly differ from the VTA_MST_ population in late juveniles (avg. 237 VTA_MST_ neurons/hemisphere vs avg. 309 VTA_MST_ neurons/hemisphere; p>0.1). These results indicate an outsized growth of the VTA_X_ projection over the same developmental period when the bird is learning its song.

## Discussion

Distinct pathways within the larger mesostriatal DA system that carry different types of reinforcement signals could enable animals to learn in complex conditions and across a wide range of objectives [1, 9, 17, 34-37]. Recent anatomical and physiological studies show that segregated pathways indeed exist and can carry distinct types of signals[7-9, 38-40]. These studies raise the question of how, and when, segregation within the mesostriatal system is formed. Past work in mammals showed that dopaminergic innervation of patch striatal areas precede matrix ones during embryogenesis [41], but it remains unclear how these results map to functionally distinct circuits that may also be spatially segregated, as in the songbird [14, 15]. Here we focused on two segregated channels in the songbird to determine the role of development and experience on DA pathway formation.

The VTA projection to the song system striatal nucleus Area X carries singing related error signals, controls striatal plasticity, and is important for song learning [16, 17, 19, 20]. A DA projection to an adjacent striatal nucleus, MST, arises from a distinct group of neurons that reside in an overlapping part of VTA, and we confirmed that the VTA_X_ and VTA_MST_ populations were distinct in adults [14, 15]. The key novel finding of the present manuscript is that this spatial segregation also exists in very young birds (<35 dph) that have had little, if any, vocal practice. These results support a model where the segregation of these pathways is genetically determined, though we cannot rule out the possibility that even earlier life experience, for example during begging calls, may play a role [42]. Future studies in even younger birds (∼10 dph) would be necessary to address this question, but due to the technical difficulties of survival neurosurgeries in bird embryos, genetic methods may need to be deployed.

We also discovered that the number of VTA_X_ neurons dramatically expands during the time course of vocal development. Past work shows that DA modulated plasticity exists in Area X during vocal development, around the same time we observe growth in the DA input to Area X [28, 29]. These data suggest that the VTA to Area X projection expands significantly during the time course of vocal development. Substantial remodeling, including invasion of axons into other song system nuclei, have also been reported during this timeframe [14, 30]. We also noted that the VTA_MST_ neuron population expands during the time course of vocal development, but the population sizes of VTA_MST_ and VTA_X_ neurons differ past 65 dph with the VTA_X_ neurons outnumbering VTA_MST_ neurons on average more than two to one in adults (Fig 1F).

If findings from the avian system generalize to mammals, our data would suggest distinct mesostriatal systems in mammals, for example that carry aversive versus reinforcing outcome signals [8, 38-40], may be patterned very early in life, even before valent experiences [41, 43]. Future studies in mammals examining mesostriatal development in pathways known to exhibit distinct types of outcome signals will be important to test for genetic versus experience-dependent modeling of the mesostriatal system.

---

**Supplementary Figure 1:**
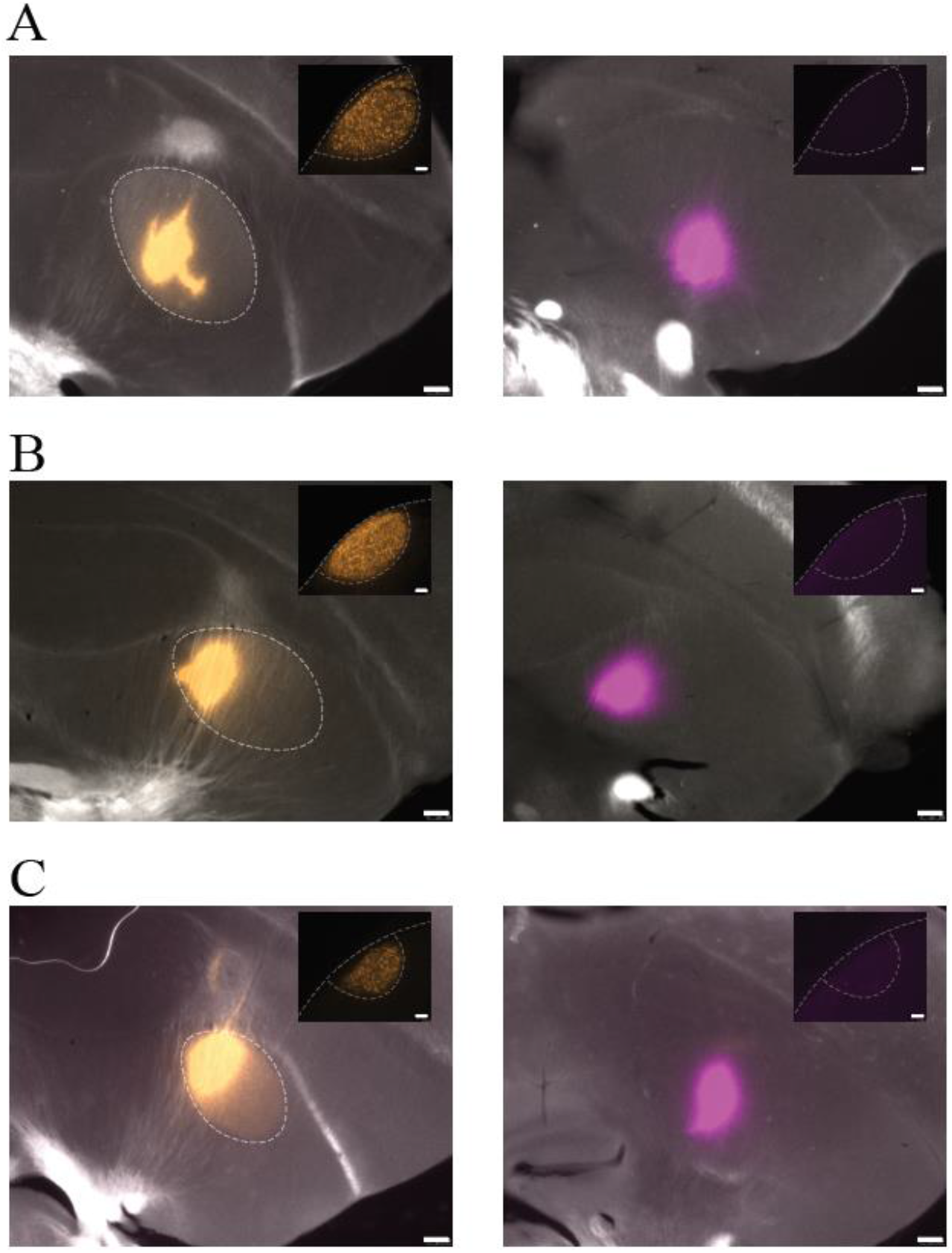
Confirmation of Area X and MST Injections. (A-C) Area X, denoted by dashed white lines, (left) and MST (right) injections. Insets show HVC, denoted by dashed white lines. Note retrogradely labeled HVC neurons following Area X injections and absence of neurons following MST injections, confirming absence of leakage from MST injections into Area X. (A) adults; (B) late juveniles; (C) early juveniles. Scale bars are 250 μm in injection images and 50 μm in inset images.

**Supplementary Table 1:** Raw Data Cell Counts for All Birds. All counted cells in VTA projecting to Area X, MST, and that are co-labeled with n = 12, n = 7, and n = 7 hemispheres for adult, late juvenile, and early juvenile injection sites respectively.

| **Bird ID** | **Injection Site** | **AreaX Projections** | **MST Projections** | **Colabeled Cells** |
| --- | --- | --- | --- | --- |
| Adult1_RH | Adult | 701 | 99 | 10 |
| Adult1_LH | Adult | 430 | 189 | 6 |
| Adult2_RH | Adult | 369 | 258 | 1 |
| Adult2_LH | Adult | 143 | 251 | 3 |
| Adult3_RH | Adult | 660 | 439 | 1 |
| Adult3_LH | Adult | 524 | 218 | 6 |
| Adult4_RH | Adult | 252 | 148 | 3 |
| Adult4_LH | Adult | 369 | 103 | 1 |
| Adult5_RH | Adult | 548 | 107 | 0 |
| Adult5_LH | Adult | 543 | 221 | 1 |
| Adult6_RH | Adult | 873 | 445 | 1 |
| Adult6_LH | Adult | 704 | 361 | 4 |
| LJuv1_LH | Late Juvenile | 48 | 194 | 0 |
| LJuv2_RH | Late Juvenile | 404 | 423 | 4 |
| LJuv2_LH | Late Juvenile | 303 | 327 | 2 |
| LJuv3_RH | Late Juvenile | 526 | 397 | 3 |
| LJuv4_LH | Late Juvenile | 311 | 312 | 4 |
| LJuv5_RH | Late Juvenile | 214 | 270 | 3 |
| LJuv5_LH | Late Juvenile | 274 | 244 | 3 |
| EJuv1_RH | Early Juvenile | 82 | 97 | 7 |
| EJuv1_LH | Early Juvenile | 79 | 79 | 3 |
| EJuv2_RH | Early Juvenile | 29 | 50 | 0 |
| EJuv2_LH | Early Juvenile | 161 | 52 | 0 |
| EJuv3_RH | Early Juvenile | 192 | 138 | 0 |
| EJuv3_LH | Early Juvenile | 112 | 90 | 0 |
| EJuv4_RH | Early Juvenile | 114 | 71 | 1 |

